# Monomeric TDP-43 binds GU-rich RNA via its NTD but only its oligomers block cryptic exon inclusion

**DOI:** 10.64898/2026.10.01.755875

**Authors:** Aurélie Zbinden, Izabela Smok, Flaminia Pellegrini, Alexandra Oertel, Antoine Cléry, Vera I. Wiersma, Wan-Ping Huang, Beatrice Gatta, Weijia Zhong, Yaozong Li, Michael Götze, Martina Peter, Niklas Bargenda, Marian Hruska-Plochan, Amedeo Caflisch, Pietro Fratta, Tatyana A. Shelkovnikova, Frédéric Allain, Alexander Leitner, Magdalini Polymenidou

## Abstract

TDP-43 aggregation is a pathological feature of several neurodegenerative diseases, resulting in nuclear loss of function and RNA dysregulation, including cryptic exon derepression. TDP-43 comprises two RNA recognition motifs required for canonical RNA binding, a low-complexity domain implicated in phase separation, and an N-terminal domain (NTD) that mediates oligomerization. While NTD-driven oligomerization is critical for TDP-43 splicing function, some studies suggest a direct interaction of NTD with RNA. Here, we investigated RNA binding by the NTD and its functional interplay with oligomerization. We show that the NTD preferentially binds UG-rich RNA via residues located at the oligomerization interface. The NTD engaged RNA only on monomeric TDP-43, as RNA binding and oligomerization compete for the same interface and rely on similar interaction modes. By disentangling these competing activities, we demonstrated that oligomerization, but not NTD-RNA interaction, is essential for TDP-43-dependent alternative splicing and repression of cryptic exons. Our findings uncover a previously unappreciated RNA-binding activity within the TDP-43 NTD that directly competes with oligomerization and establish oligomerization as a key determinant of TDP-43 cryptic exon repression.

## Introduction

TAR DNA-binding protein 43 (TDP-43) is a predominantly nuclear protein, that undergoes cytoplasmic mislocalization and aggregation, a defining pathological hallmark of both amyotrophic lateral sclerosis (ALS) and frontotemporal dementia (FTD), two fatal and incurable neurodegenerative diseases (Arai et al., 2006; Ling et al., 2013; Neumann et al., 2006). TDP-43 is an RNA-binding protein (RBP) that plays a central role in RNA homeostasis, regulating transcripts essential for neuronal health and integrity (Brown et al., 2022; El-Agamy et al., 2026; Joseph et al., 2025; Keuss et al., 2024; Klim et al., 2019; Melamed et al., 2019; Polymenidou et al., 2011; Tollervey et al., 2011). Among its best characterized functions are the regulation of alternative splicing and the repression of cryptic exons (CE), processes mediated through binding to GU-rich intronic regions (Brown et al., 2022; El-Agamy et al., 2026; Ling et al., 2015; Melamed et al., 2019; Polymenidou et al., 2011; Tollervey et al., 2011). Consistent with these roles, loss of nuclear TDP-43 leads to widespread transcriptome dysregulation, resulting in aberrant isoforms and altered gene expression levels.

Considerable attention has focused on the role of TDP-43 in repressing CE. Upon loss of nuclear TDP-43, cryptic exons are aberrantly included in transcripts, often resulting in frameshifts and the introduction of premature stop codons that trigger nonsense-mediated decay, ultimately leading to loss of functional protein (Brown et al., 2022; El-Agamy et al., 2026; Ling et al., 2015). This emphasis on CE is supported by findings showing that CE inclusions occur specifically in ALS–FTD patient, but not control, brains (Brown et al., 2025, 2022; Ling et al., 2015; Prudencio et al., 2020). Importantly, many of the affected transcripts are essential for neuronal function, some exhibit neuron-specific expression (e.g., STMN2), and key events such as CE inclusion in UNC13A have been associated with disease progression (Brown et al., 2025, 2022; El-Agamy et al., 2026; Keuss et al., 2024; Klim et al., 2019; Melamed et al., 2019).

TDP-43 belongs to the heterogeneous nuclear ribonucleoprotein (hnRNP) family and consists of 414 amino acids organized into four functional domains (Afroz et al., 2017; Buratti et al., 2001; Conicella et al., 2016; Lukavsky et al., 2013; Zbinden et al., 2020). These include two RNA recognition motifs (RRMs, aa 106–262), responsible for binding GU-rich RNA sequences (Buratti et al., 2001; Lukavsky et al., 2013); and a C-terminal intrinsically disordered low-complexity domain (LCD, aa 263–414), essential for TDP-43 liquid-liquid phase separation (LLPS) and protein-protein interactions (Buratti et al., 2005; Conicella et al., 2016; Feneberg et al., 2020) and an N-terminal domain (NTD, aa 1–80), which mediates self oligomerization (Afroz et al., 2017, 2017; Chang et al., 2012; Shiina et al., 2010).

TDP-43 self-oligomerization is dynamic and concentration-dependent, and is mediated by head-to-tail interactions between two NTDs (Afroz et al., 2017). Specifically, the negatively charged tail region (E14–Q34) interacts with the positively charged head region (Q49–L56), primarily through electrostatic interactions (Afroz et al., 2017). These resulting dimers and higher-order oligomers are predominantly localized in the nucleus, although smaller populations have also been reported in the cytoplasm (Afroz et al., 2017; Pérez-Berlanga et al., 2023). We and others have demonstrated that physiological TDP-43 oligomerization is critical for multiple aspects of its function, including LLPS, splicing regulation, autoregulation, and nuclear retention, and protects against cytoplasmic aggregation (Afroz et al., 2017; Jiang et al., 2017; Koehler et al., 2022; Oiwa et al., 2023; Pérez-Berlanga et al., 2023; A. Wang et al., 2018; Yu et al., 2021; Zbinden et al., 2020; Zhang et al., 2013).

Notably, several studies have also suggested a direct interaction of the NTD with nucleic acids, indicating that this domain may contribute to both oligomerization and RNA binding. Indeed, the folding equilibrium of the NTD can be shifted to its native state in the presence of (TG)_6_ single-stranded DNA (ssDNA) (Qin et al., 2014), which directly binds purified NTD in electrophoretic mobility shift assays (EMSA) (Afroz et al., 2017). Moreover, the presence of TT and TG repeats, but not random ssDNA, induces the NTD to undergo LLPS in a sequence- and concentration-dependent manner (Wang et al., 2018). While these studies provide indications for direct nucleic acid interaction with the NTD, the exact binding mode, binding mechanism, sequence specificity and its interplay with oligomerization remain elusive.

Here, we sought to investigate NTD-mediated RNA binding and its interplay with oligomerization in regulating TDP-43 function. We identify a clear RNA-binding site within the NTD oligomerization interface and demonstrate that this interaction occurs exclusively in the monomeric state of the protein. Our structural analysis reveals that this exclusivity arises because RNA binding and oligomerization utilize the same interaction interface and rely on similar interaction types. By disentangling these competing activities, we demonstrate that oligomerization, but not NTD-RNA interaction, is essential for TDP-43-dependent alternative splicing and efficient repression of cryptic exons.

## Results

### The NTD of TDP-43 interacts with GU-repeat RNA within its oligomerization interface

Over the past decade, several studies have suggested that the behavior and structure of the NTD of TDP-43 are affected by the presence of nucleic acids (Afroz et al., 2017; Qin et al., 2014; Wang et al., 2018). To elucidate the existence of RNA binding sites within the NTD, we used cross-linking of isotope-labeled RNA and subsequent tandem mass spectrometry (CLIR-MS), a method that allows high-resolution mapping of interactions in RNA-protein complexes on both the protein and RNA level (Dorn et al., 2017; Götze et al., 2021) (Figure 1A). We purified several recombinant TDP-43 protein fragments: the NTD alone (NTD: amino acids (aa) 1-80), a construct containing both RNA recognition motifs (RRMs) RRM1 and RRM2 (RRMs^WT^: aa 106-262), an RNA-binding–deficient RRM mutant (RRMs^mut^: F147A/F149A/F194A/F229A/F231A) that preserves the structural integrity of the RRMs (Pérez-Berlanga et al., 2023) and a construct containing both the NTD and RRMs but lacks the low-complexity domain (NTD-RRMs: aa 1-263) (Supplementary Figure 1A and Supplementary Figure 1B). Upon cross-linking wild-type (WT) recombinant proteins to UG/GU-rich RNA (Lukavsky et al., 2013), the preferred binding motif of TDP-43 (Polymenidou et al 2011; Tollervey et al, 2011), we detected RNA interactions in the RRMs (Figure 1B-C), as expected. Cross-links within the RRMs revealed a prominent interaction site in RRM2, centered around Phe194 (K192–G196), a region critical for TDP-43 RNA binding and splicing (Lukavsky et al., 2013). This interaction was significantly diminished in our RRMs^mut^ as counts per cross-link site went drastically down confirming specificity of our experimental setup (Figure 1B, Supplementary Figure 1C). Surprisingly, we also detected a prominent interaction site in both the isolated NTD and the NTD-RRMs, with cross-links assigned to Ser48, Gln49, Cys50, and Arg52 (Figure 1B-C). Structural mapping of these residues visualized an RNA interaction site within the NTD oligomerization interface, specifically in its positively charged head region, which is essential for NTD-mediated oligomerization (Figure 1D). Given the importance of NTD-mediated oligomerization in regulating both TDP-43 physiological functions (Afroz et al., 2017; Jiang et al., 2017; Koehler et al., 2022; Oiwa et al., 2023; Pérez-Berlanga et al., 2023; Wang et al., 2018; Zhang et al., 2013) and its transition to pathology (Afroz et al., 2017; Jiang et al., 2017; Oiwa et al., 2023; Pérez-Berlanga et al., 2023; Yu et al., 2021; Zbinden et al., 2020; Zhang et al., 2013), this RNA binding directly within the oligomerization interface is likely to have important consequences for protein behavior.

**Figure 1:**
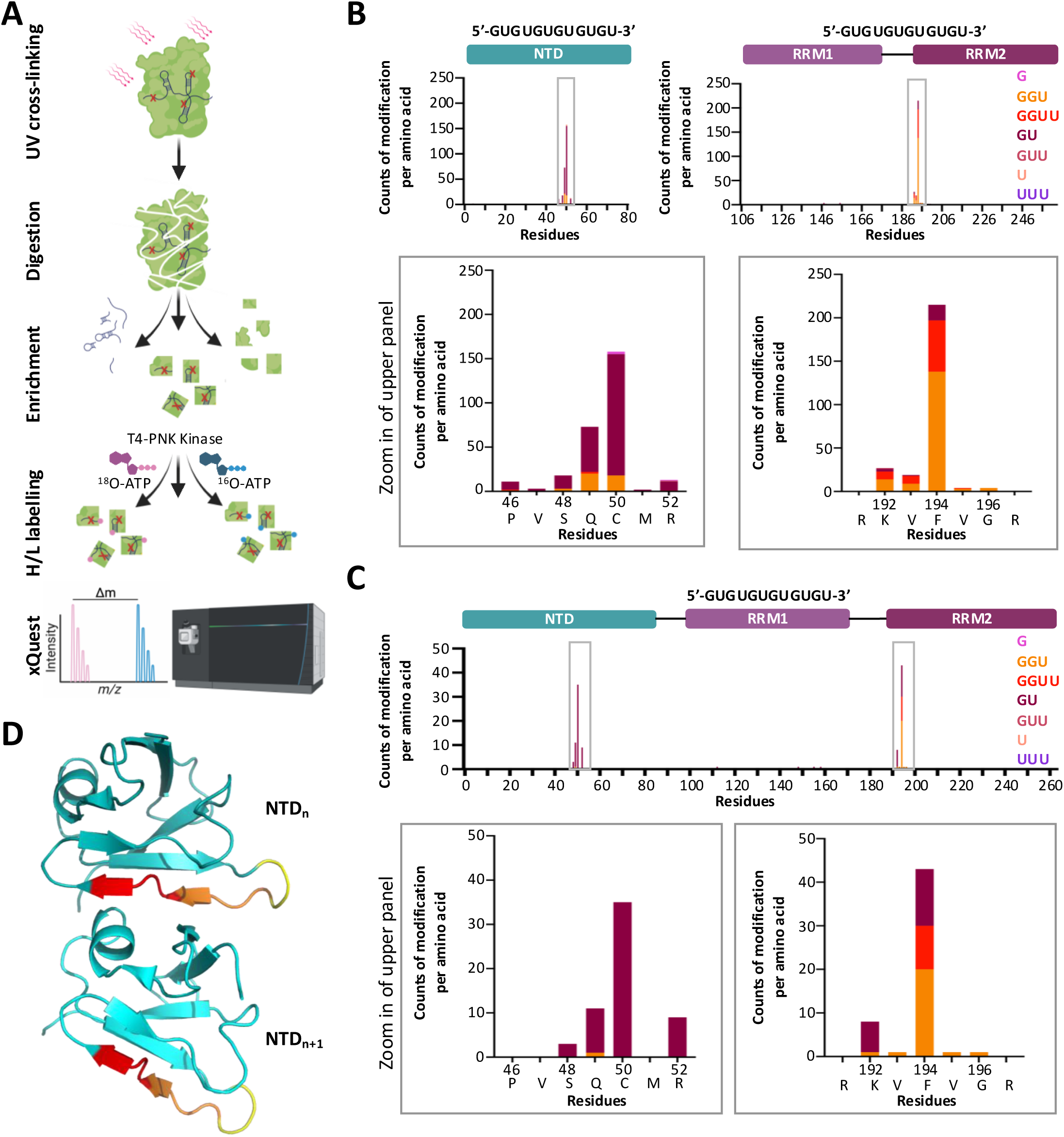
The NTD of TDP-43 interacts with RNA within its oligomerization interface. **A** Schematic representation of **C**ross-**L**inking of **I**sotope-labeled **R**NA and tandem **M**ass **S**pectrometry (**CLIR-MS**) workflow for high-resolution RNA-protein interaction mapping (Dorn et al., 2017; Götze et al., 2021; Sarnowski et al., 2023). **B** Identified (GU)_6_ RNA–protein CLIR-MS cross-links in the N-terminal domain (NTD) only (left), the RNA recognition motif (RRM) construct containing both RNA recognition motif 1 (RRM1) and RNA recognition motif 2 (RRM2) (right). Cross-links are displayed according to sequence position and shown with color-coded nucleic acid composition as indicated in the legend. Counts represent cross-links identified by spectral matches. Schematic representations of the sequence space of each construct are shown at the top. Lower panels depict zoomed-in views of the cross-linking sites shown above, with sites labelled by number and corresponding amino acid. Cross-linking was performed at a 1:1 molar ratio of protein:RNA using a protein concentration of 50 µM. **C** Identified (GU)_6_ RNA–protein CLIR-MS cross-links in the delta low-complexity domain (NTD-RRMs) construct containing the NTD and RRMs. Cross-links are displayed according to sequence position and shown with color-coded nucleic acid composition as indicated in the legend. Counts represent cross-links identified by spectral matches. Schematic representations of the sequence space of each construct are shown at the top. Lower panels depict zoomed-in views of the cross-linking sites shown above, with sites labelled by number and corresponding amino acid. Cross-linking was performed at a 1:1 molar ratio of protein:RNA using a protein concentration of 50 µM. **D** Structural representation of the interaction site identified by CLIR-MS (yellow) in comparison with previously defined NTD head region (red; (Afroz et al., 2017)). The overlapping residues are shown in orange (PDB 5mdi). NTD_n_ and NTD_n+1_ represent two NTD in the dimer constellation.

### NTD-mediated RNA binding is exclusive to the monomeric state of TDP-43

Since the NTD-mediated RNA binding mapped to the oligomerization interface, we sought to determine how oligomerization influences this interaction. To this end, we performed molecular dynamics (MD) simulations to assess UG-RNA binding in both the monomeric and oligomeric states of the NTD. In the monomeric state, the NTD exhibited RNA interaction patterns consistent with those previously identified by CLIR-MS (Figure 2A; Supplementary Figure 2A). In contrast, these interactions were largely abolished in the dimeric state, indicating that oligomerization occludes the RNA-binding interface (Figure 2A). Intriguingly, little to no interaction was detected in the surface-exposed head region of the second NTD either (Figure 2A), potentially due to the occurrence of structural rearrangements upon NTD dimerization, as recently reported (Herrera et al., 2023).

**Figure 2:**
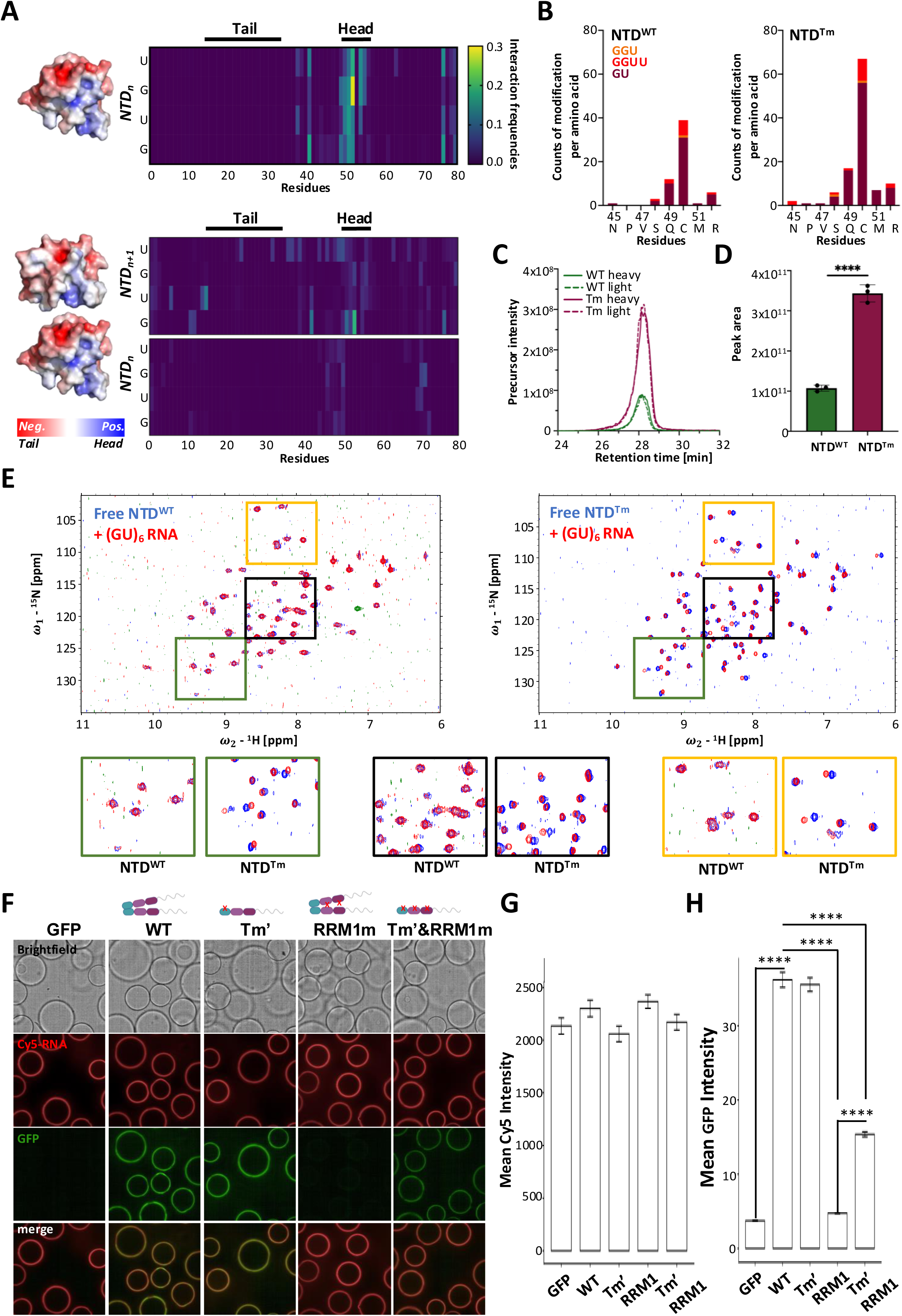
NTD-mediated RNA binding is exclusive to the monomeric state of TDP-43. **A** On the left: Monomeric (NTD_n_, top) and dimeric (NTD_n_ and NTD_n+1_, bottom) TDP-43 NTDs shown as surface charge structures (PDB 5mdi). Negative surfaces are shown in red, corresponding to the tail region of the domain, while the positive surface of the head is shown in blue. Charge legend depicted below. On the right: contact heat maps showing the interaction frequencies of the sequence space of monomeric (top) and dimeric (bottom) TDP-43 with (GU)_2_ RNA generated by molecular dynamics (MD) simulations. Tail and head regions are annotated above the graph. **B** Direct comparison of (GU)_6_ RNA–protein cross-links in TDP-43 NTD^WT^ (left) or TDP-43 NTD^Tm^ (right) identified by conventional CLIR-MS approach, shown in the sequence position and with color-coded nucleic acid composition as indicated by the legend. Counts represent cross-links identified by spectral matches. Cross-linking was performed at a molar ratio of 1:1 protein:RNA. Protein concentration was 50 µM. **C** Representative elution profile of selected cross-linked products [GU-Y_43_RNPVSQCMR_52_ (3+ precursor)] of TDP-43 NTD^WT^ (green) and TDP-43 NTD^Tm^ (burgundy) in a targeted CLIR-MS experiment. Both heavy (solid line) and light (dotted line) labeled variants are shown. Cross-linking was performed at a molar ratio of 1:1 protein:RNA. Protein concentration was 50 µM. **D** Quantification of area under the curve (AUC) by integration of elution profiles shown in (C). N=3 independent experiments. Graph bars represent mean ± SD. Unpaired two-tailed t-test was used for statistical analysis. **** p < 0.0001. **E** Overlay of 2D ^1^H-^15^N HSQC spectra of ^15^N isotope labeled TDP-43 NTD^WT^ (left) and TDP-43 NTD^Tm^ (right) in free form (blue) or with (GU)_6_ RNA at a protein:RNA ratio of 1:4 (red). Below are enlarged views of different spectral regions, color-coded in green, black, and yellow to allow visual comparison of chemical shift perturbations (CSPs) between the two NTDs. Protein concentration used was 100 µM. **F** Representative brightfield and fluorescence images of confocal nanoscanning (CONA) assay assessing RNA binding capacity of different GFP-TDP-43 variants in cell lysates. Beads were coated with Cy5-labelled (GU)_15_ RNA (synthetic oligonucleotide) and were incubated with lysates from cells expressing GFP-tagged TDP-43 variants, including the wild-type (WT) TDP-43, monomeric Tm’ variant (E14A/E17A/E21A), an RRM1 mutant (F147A/F149A) and a variant combining both Tm’ and RRM1 (E14A/E17A/E21A/F147A/F149A). **G** Mean GFP ring fluorescence (F) was quantified. A total of 95–160 beads per condition were analyzed using an automated pipeline (6–10 beads per field of view; 17 fields total). **** p < 0.0001, one-way ANOVA with Tukey’s post hoc test. Normalized mean Cy5 fluorescence across all images confirming comparable coating of (GU)_15_ RNA across all conditions is also shown.

To validate our MD simulations, we compared the wild-type NTD (NTD^WT^), which exists in a concentration-dependent equilibrium of monomers, dimers, and higher-order oligomers, with a constitutively monomeric variant, the NTD tail mutant (NTD^Tm^). The previously described NTD^Tm^ harbors two mutations (E17A/E21A) in the tail part of the NTD and has been demonstrated to successfully abolish oligomerization *in vitro* without affecting the structural integrity of the domain (Afroz et al., 2017). Since the mutations are far away from the head region, which interacts with RNA, we reasoned that this mutant would ensure the preservation of the RNA binding by the NTD while neutralizing oligomerization (Supplementary Figure 2B). We first performed conventional CLIR-MS of NTD^WT^ and NTD^Tm^ with a GU-rich RNA to compare the RNA interaction propensity of both states. Indeed, monomeric NTD^Tm^ showed an increase in detected cross-links compared to the NTD^WT^ (Figure 2B; Supplementary Figure 2C). While conventional CLIR-MS serves as a robust tool for localizing RNA binding regions, it possesses certain limitations, including a semi-quantitative nature due to the use of spectral counting. To overcome these constraints, we used a targeted MS approach based on parallel reaction monitoring (PRM), enabling the quantification of predefined targets of interest between multiple variants (Supplementary Figure 2D). Using this targeted approach with the peptide-RNA cross-link GU-Y_43_RNPVSQCMR_52_, covering the head part of the oligomerization interface, we saw that the interaction to GU-RNA was significantly increased in the monomeric NTD^Tm^ compared to NTD^WT^ (Figure 2C-D, Supplementary Figure 2G).

To validate this finding, we purified ^15^N-labeled NTD^WT^ and NTD^Tm^ (Supplementary Figure 2F) and recorded heteronuclear single quantum coherence (^1^H-^15^N HSQC) nuclear magnetic resonance (NMR) spectra (Figure 2E). In line with our previous data (Afroz et al., 2017), the structural integrity of the NTD was not compromised by the monomerizing Tm mutations and observed differences in spectral coverage with the NTD^WT^ were due to the amino acid substitutions and oligomerization (Supplementary Figure 2G). In addition, the NTD^WT^ exhibited broader peaks characteristic for interaction with medium exchange rates, demonstrating that the NTD^WT^, but not the NTD^Tm^, undergoes oligomerization under these conditions and represents the dominant state observable by NMR (Supplementary Figure 2G). When GU-rich RNA was titrated to either the NTD^WT^ or monomeric NTD^Tm^ only the latter showed chemical shift perturbations (CSPs) verifying that only the monomeric form can interact with RNA (Figure 2E).

The absence of NMR-detectable RNA binding in NTD^WT^, in contrast to our mass spectrometry approach, likely reflects the co-existence of multiple oligomeric states in the NTD^WT^ (monomeric, dimeric and multimeric) and the different underlying principles of the biophysical techniques used. While CLIR-MS approach allows for the detection of fragments from all NTD states, including the monomer, NMR as an ensemble technique only reflects the most dominant state, in this case the multimeric RNA-unbound NTD (Supplementary Figure 2G). To assess if monomeric NTD-mediated RNA binding also occurs in the full-length protein under more physiological conditions we utilized a confocal nanoscanning (CONA) assay (Zhu et al., 2021). In short, beads coated with a Cy5-labelled GU-rich RNA were incubated with cell lysate from cells expressing full-length GFP-tagged TDP-43 variants, allowing us to assess RNA binding propensity in the complex cell lysate environment (Supplementary Figure 2H). We compared WT GFP-TDP-43 with a monomeric mutant (Tm′; E14A/E17A/E21A), an RNA-binding–deficient RRM1 mutant (RRM1m; F147A/F149A), and the corresponding double mutant (Tm′ RRM1m). As expected, both WT and Tm′ exhibited robust binding, whereas the GFP-only control and the RRM1 mutant showed no detectable interaction (Figure 2F-H). Notably, the monomeric double mutant (Tm′ RRM1m) retained RNA binding albeit less than WT and Tm, indicating that loss of canonical RRM-mediated binding can be compensated by the NTD, but only when TDP-43 is in a monomeric state (Figure 2F-H). Taken together, these results provide compelling evidence that NTD-mediated RNA binding occurs in the monomeric state of TDP-43.

### A shared interaction interface governs both NTD-RNA binding and oligomerization

To understand the apparently preferential RNA interaction to the monomeric state of TDP-43 we performed three-dimensional (3D) NMR with ^13^C-^15^N-labeled monomeric NTD^Tm^ to assign the backbone amides of the protein (Figure 3A, Supplementary Figure 3A-B). We discovered 33 amino acids shifting with a CSP greater than or equal to 0.02 in the presence of GU-rich RNA (Figure 3A and Supplementary Figure 3B). Most dominant CSPs (≥ 0.04) were observed for Leu41, Tyr43, Asn45, Ser48, Gln49, Arg52, Gly53, Val54, Arg55, Leu56, Val57, Glu58, Gly59, His62, Asn76, Tyr77, Asp80 and Asn81 most of which are evolutionary conserved residues (Supplementary Figure 3C). When mapped on the structure, these residues delineated an interaction surface within the oligomerization interface, predominantly concentrated in the head region (Gln49-Leu56) validating the previous findings from our CLIR-MS and MD simulation approaches (Figure 3B-C). When comparing the residues involved in RNA binding (CSP ≥ 0.04) and oligomerization (Afroz et al., 2017), 9 out of the 17 amino acids identified were important for both oligomerization and RNA binding (Figure 3D). Mapping of these residues responsible for RNA binding alone and those contributing to both RNA binding and oligomerization on the structure showed both sets occupying the same spatial region, suggesting structural overlap for both functions (Figure 3E, Supplementary Movie 1).

**Figure 3:**
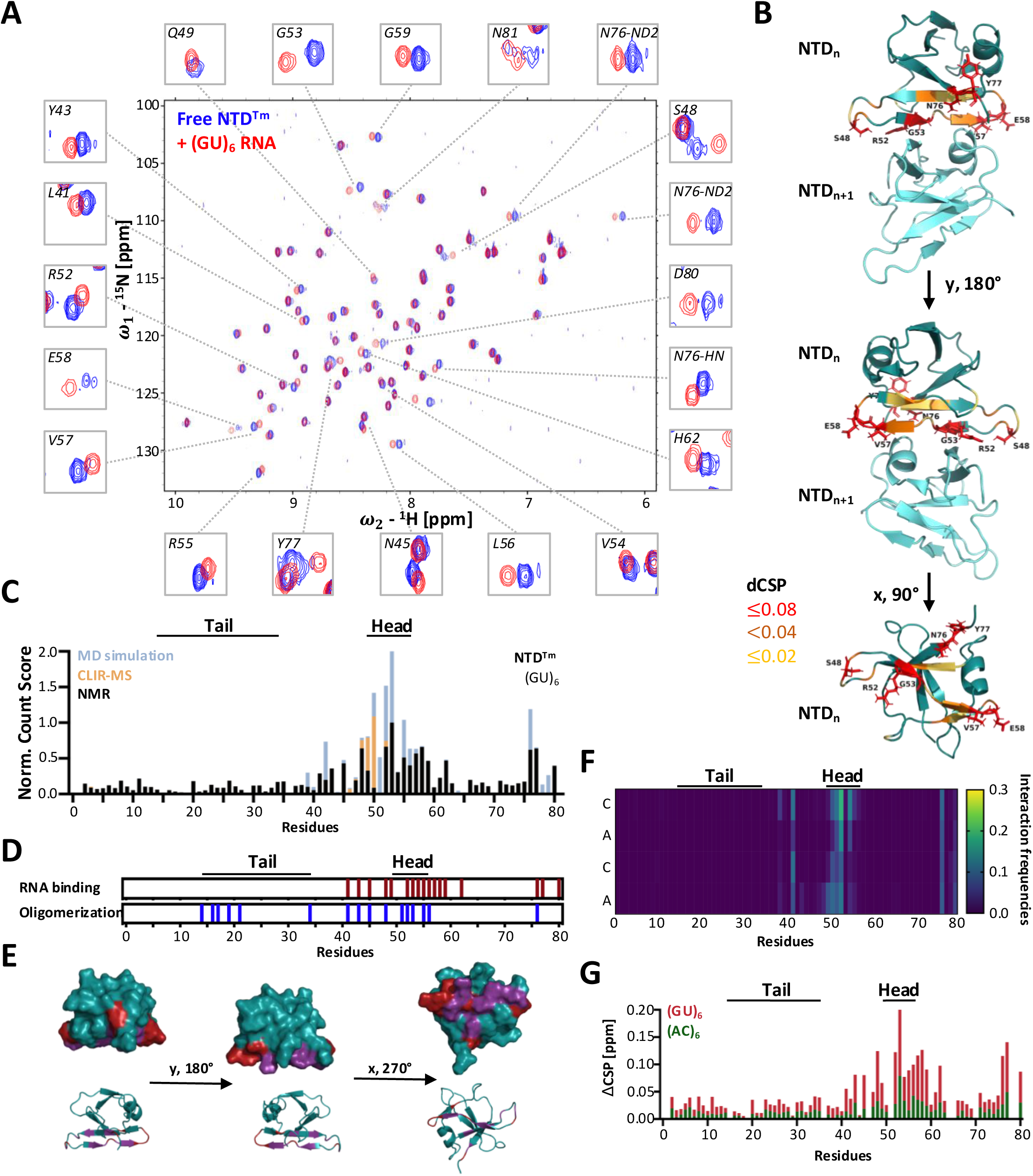
A shared interaction interface governs both NTD-RNA binding and oligomerization. **A** Overlay of 2D ^1^H-^15^N HSQC spectra of ^15^N isotopically labeled free TDP-43 NTD^Tm^ (blue) and saturated with (GU)_6_ RNA (red). Magnified views of cross peaks with chemical shift perturbations are shown around the full spectra and are annotated with the corresponding assignments elucidated by 3D ^1^H-^15^N-^13^C NMR. NTD^Tm^ concentration used was 100 µM with a protein:RNA molar ratio of 1:8. **B** Cartoon representation of NTD dimer (PDB 5mdi) with highlighted amino acid backbones based on calculated average chemical shift perturbations ΔCSP = ((ΔδH)^2 + (ΔδN/6.51)^2)^(1/2) from the free and saturated (GU)_6_ signal. Different colors represent different strengths of CSP as indicated by the legend on the right. Amino acids with a CSP equal or greater than 0.08 are shown in stick representation from different angles. **C** Normalized (Norm.) count score of TDP-43 NTD^Tm^ amino acids with (GU)_6_ RNA from NMR, CLIR-MS experiments, and molecular dynamics simulations to visualize the overall fit of the identified RNA interaction regions generated by different biophysical methods. NMR is represented by calculated per amino acid backbone amide average ΔCSP = ((ΔδH)^2 + (ΔδN/6.51)^2)^(1/2) between the free and (GU)_6_ RNA-bound form. CLIR-MS signal represents the sum of counts of all nucleic acids cross-linked to each amino acid identified by conventional approach. MD signal represents the sum of all contact frequencies assigned to each amino acid of the monomeric MD simulation. Data from each experimental approach have been normalized (values between 0 and 1) to circumvent the issue of incomparable intensities across the different biophysical methods. **D** Comparison between amino acids important for oligomerization (blue, based on previously published work (Afroz et al., 2017) and amino acids important for NTD-mediated (GU)_6_ RNA binding based on NMR measurements (burgundy) over the sequence space of the NTD of TDP-43. **E** Surface (top) and cartoon (bottom) views of monomeric NTD shown from different rotation angles (PDB 5mdi). Amino acid backbones highlighted in purple are involved in both oligomerization and NTD-mediated (GU)_6_ RNA binding, while those in red are only involved in RNA binding. **F** Heat map showing interaction frequencies over the sequence space of the monomeric TDP-43 with (AC)_2_ RNA generated by molecular dynamics simulations. Tail and head regions are annotated above the graph. **G** Plotted chemical shift perturbations (ΔCSP) = ((ΔδH)^2 + (ΔδN/6.51)^2)^(1/2) of saturated (AC)_6_ RNA in green or (GU)_6_ RNA in red as a stacked bar plot representation over the sequence space of the NTD, with annotated tail and head regions above. ΔCSP were generated relative to the free form of the monomeric NTD^Tm^ at 100 µM protein concentration.

Oligomerization is predominantly driven by electrostatic interactions, including the contribution of two positively charged arginines in the head regions (R52 and R55) (Afroz et al., 2017). To assess the contribution of such interactions to NTD-mediated RNA binding, we mutated aforementioned arginine residues (head mutant (Hm); R52 and R55), which we also identified by NMR as key contributors to RNA interaction (Figure 3A, Supplementary Figure 3D). ^1^H-^15^N HSQC NMR of ^15^N-labeled NTD^Hm^ showed no compromise in the structural integrity of the NTD^Hm^ and differences in spectral coverage with the NTD^WT^ can be attributed to amino acid mutations and oligomerization (Supplementary Figure 3E-F). The addition of GU-rich RNA did not lead to any observable CSP, peak alteration, or peak disappearances, suggesting the absence of significant RNA binding by NTD^Hm^ (Supplementary Figure 3G).

Collectively, these findings indicate that NTD-mediated RNA binding is restricted to the monomeric state, as it competes directly with oligomerization for the same residues, structural interface, and electrostatic interactions.

### TDP-43 NTD preferentially binds to GU-rich RNA

TDP-43 binds numerous transcripts enriched in GU repeats (Polymenidou et al., 2011; Tollervey et al., 2011), consistent with the GU-sequence specificity of the RRMs (Lukavsky et al., 2013). To assess the sequence specificity of monomeric NTD RNA binding, we performed CLIR-MS experiments with AC-repeat RNA and several of our recombinant TDP-43 protein fragments (Supplementary Figure 4A-B, Supplementary Figure 1A-B). Compared to GU-repeats (Figure 1B-C), cross-linking to AC-repeats was drastically reduced for the RRMs and NTD^WT^. Furthermore, we observed no AC-repeat cross-links to the NTD and only minimal cross-links to the RRMs in the NTD-RRMs TDP-43 variant (Figure 1B-C, Supplementary Figure 4A-B). In agreement, the MD simulation of monomeric NTD with AC-rich RNA leads to significantly lower interaction frequencies than with GU-rich RNA (Figure 2A, Figure 3F). To validate these findings, we performed NMR with ^15^N-labeled NTD^WT^, NTD^Hm^ and NTD^Tm^ in the presence of AC- or GU-rich RNA (Figure 3G, Supplementary Figure 4C). We did not observe any interactions between NTD^WT^ or NTD^Hm^ and AC-rich RNA (Supplementary Figure 4C), while NTD^Tm^ showed dramatically diminished interactions with AC-rich RNA, compared to GU-rich RNA (Figure 3G). Only 3 out of the 16 identified GU-RNA interacting amino acids showed a ΔCSP of ≥ 0.04 in the presence of AC-rich RNA (Gly53, Leu56, Tyr77) in line with an RNA sequence preference for GU-repeats.

To further validate the difference in GU- and AC-RNA binding by the monomeric NTD^Tm^ we conducted isothermal calorimetry (ITC) experiments. We could determine an approximate dissociation constant (K_D_) of 190 μM +/- 55 for the NTD^Tm^ to GU-rich RNA (Supplementary Figure 4D) with a N value of 7.5 +/- 2. In contrast, our attempts to perform ITC with NTD^Tm^ and AC-rich RNA failed to generate a binding isotherm, rendering it impossible to determine a proper K_D_ value for this interaction (Supplementary Figure 4D), consistent with an extremely weak interaction. Collectively, our findings show that the monomeric NTD exhibits a preference for binding to GU-containing RNAs.

### Cryptic exon repression relies solely on oligomerization and not NTD-mediated RNA binding

To investigate NTD-mediated RNA binding and its functional interplay with oligomerization in TDP-43-dependent splicing, we integrated the TDP-REG reporter, which is based on mScarlet protein expression regulated by the inclusion of a TDP-43-dependent cryptic exon (CE), within HeLa TDP-43 knockout (KO) cells (Roczniak-Ferguson and Ferguson, 2019). The latter are thoroughly characterized cells with quantifiable morphological changes that can be rescued by re-introduction of wild-type (WT) TDP-43 (Roczniak-Ferguson and Ferguson, 2019). The resulting HeLa KO cells stably expressing the TDP-REG reporter showed constant mScarlet protein expression due to the loss of TDP-43 (Figure 4A). To determine the functionality of TDP-43 variants in CE repression, we transiently transduced HeLa TDP-43 KO reporter cells with the following GFP-tagged TDP-43 variants: GFP-TDP-43-WT, monomeric Tm mutant retaining the NTD-mediated RNA binding (GFP-TDP-43-Tm), monomeric Hm mutant lacking the NTD-mediated RNA binding (GFP-TDP-43-Hm), and the well-characterized monomeric 6M variant (Afroz et al., 2017; Pérez-Berlanga et al., 2023), combining both the Tm and Hm mutations along with two additional amino acid substitutions (GFP-TDP-43-6M) (Supplementary Figure 5A). We used fluorescence-activated cell sorting (FACS) analysis of cells expressing equal levels of the GFP-TDP-43 variants, thereby minimizing dose-dependent effects (Figure 4B). Notably, in contrast to GFP-TDP-43-WT-expressing cells, which lost mScarlet expression due to restoration of CE splicing repression, sorted cells with equal expression of all monomeric mutants (Tm, Hm, 6M) failed to rescue the reporter activity, indicating that oligomerization is necessary for CE repression (Figure 4B-C; Supplementary Figure 5B-C).

**Figure 4:**
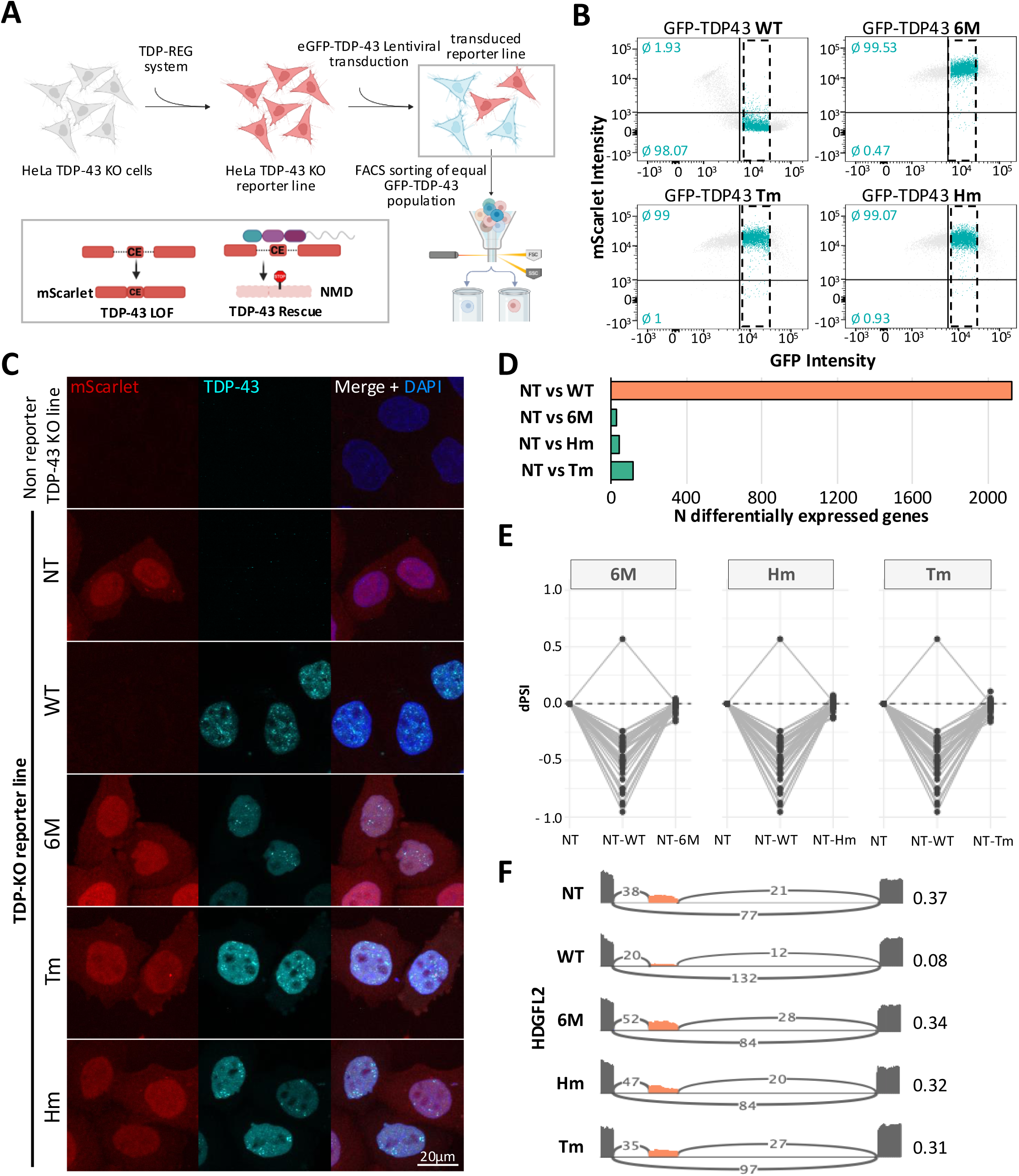
Cryptic exon repression relies solely on oligomerization and not NTD-mediated RNA binding. **A** Schematic representation of HeLa knock-out (KO) TDP-43-REG reporter lines used for RNA sequencing analysis. Reporter lines are transduced with GFP-tagged TDP-43 variants, sorted by FACS and gated using the GFP signal to achieve equal TDP-43 levels among different variants to avoid concentration-dependent effects. Below: Schematic representation of the TDP-REG mScarlet reporter (Wilkins et al., 2024). In the absence of TDP-43 KO, the cryptic exon (CE) is included, resulting in the expression of the red mScarlet reporter. Upon TDP-43 reintroduction (rescue), CE inclusion is suppressed, causing a frameshift that triggers nonsense-mediated decay (NMD) and silencing of mScarlet expression. **B** Representative FACS scatter plots after 6 days of transduction showing GFP and mScarlet intensity distribution among different GFP-TDP variants. Dotted boxes indicate selected cell population for further processing. Numbers annotated in light blue depict the average percentage (N=3) of cells with mScarlet positivity (upper number) or negativity (lower number). **C** Representative immunofluorescence images of cells collected by FACS expressing different GFP-TDP-43 variants. mScarlet (red) is silenced upon restoration of correct splicing by wild-type GFP-TDP-43-WT. The original, non-reporter TDP-43 KO cell line is included as a control. TDP-43 is shown in turquoise, nuclei are stained with DAPI (blue). Scale bar: 20 µm. **D** Differential expression analysis comparison between non-transduced (NT) and GFP-TDP-43 variants shows clear rescue of gene expression only in the GFP-TDP-43-WT. **E** Differential percentage spliced in analysis (dPSI) of known cryptic events (Brown et al., 2025) comparing NT, NT-WT, and NT to each monomeric variant, showing the same trend across all variants. **F** Representative sashimi plots of the cryptic exon (orange) coverage in the HDGLF2 gene. PSI values for each variant are annotated on the right.

To assess how oligomerization and NTD-mediated RNA binding impact TDP-43 function at the transcriptome level, we performed RNA sequencing (RNA-seq) on transduced HeLa TDP-43 KO reporter cells sorted based on equal expression of GFP-TDP-43 variants (Figure 4B). At comparable expression levels of all GFP-TDP-43 variants (Supplementary Figure 5D-E), the number of differentially expressed genes compared to the non-transduced (NT) KO cells was significantly lower for all monomeric variants in contrast to GFP-TDP-43-WT-expressing cells (Figure 4D). Specifically, we observed 2128 differentially expressed genes upon GFP-TDP-43-WT expression compared to NT HeLa KO cells, while expression of all monomeric variants resulted in minimal changes in gene expression, i.e. only 32 (6M), 47 (Hm), and 116 (Tm) genes were differentially expressed (Figure 4D, Supplementary Figure 5F). Hierarchical clustering analysis confirmed that all monomeric mutants clustered with the non-transduced (NT) condition, whereas GFP-TDP-43-WT formed a distinct cluster (Supplementary Fig 5G). Consistently, genes differentially expressed upon GFP-TDP-43-WT expression showed an anticorrelation not only with NT cells, but also with all three monomeric TDP-43 variants, which instead displayed a positive correlation with the NT condition (Supplementary Figure 5H). In summary, we did not observe significant differences in gene expression among monomeric variants, irrespective of their ability to bind RNA via the NTD (Tm versus Hm and 6M).

TDP-43-dependent transcriptomic changes are widespread and may include indirect effects, as TDP-43 regulates the expression of other RNA-binding proteins (RBPs), which can in turn alter gene expression (Polymenidou et al., 2011; Tollervey et al., 2011). In contrast, cryptic exon inclusion represents a more direct and specific readout of TDP-43 loss of function (Brown et al., 2025, 2022; El-Agamy et al., 2026). We therefore investigated whether differences between monomeric variants lead to differences in cryptic splicing events. All cryptic junctions that were significantly suppressed in GFP-TDP-43-WT-expressing cells compared to the NT condition remained unchanged in all three monomeric mutants (Figure 4E; Supplementary Figure 5I–L), indicating that monomeric TDP-43 cannot rescue cryptic repression. This included a well-characterized cryptic exon event in the *HDGFL2* gene (Figure 4F, Supplementary Figure 5M), a cryptic junction specifically observed in ALS and FTD patient brains (Albagli et al., 2024; Irwin et al., 2024; Seddighi et al., n.d.). Together, these results indicate that monomeric TDP-43 variants are unable to restore canonical splicing regulation, irrespective of their capacity for NTD-mediated RNA binding, and that splicing regulation, including CE repression, necessitates NTD-driven oligomerization.

## Discussion

In this study, we sought to determine whether the NTD of TDP-43 harbors an RNA-binding site and how this interaction functionally interplays with oligomerization in splicing regulation. We identified a previously unrecognized RNA-binding site within the positively charged head region (Q49–L56), which exhibits preferential binding to GU-rich sequences. Importantly, we showed that this interaction occurs exclusively in the monomeric state. Surprisingly, the NTD can partially compensate for loss of RRM1-mediated RNA binding, but only under monomeric conditions. Structural analysis further revealed that RNA binding and oligomerization occupy the same interface, involve overlapping residues, and rely on similar electrostatic interactions. When studying the functional interplay between oligomerization and NTD-mediated RNA binding we elucidated that for splicing regulation, including cryptic exon repression, TDP-43 oligomerization is essential.

The NTD shares strong structural similarity with the polymer-forming Dishevelled and Axin (DIX) domains (Afroz et al., 2017; Qin et al., 2014). While such domains are well known for their dynamic head-to-tail oligomerization, they are not generally associated with nucleic acid binding (Bienz, 2014). Although previous studies have hinted at this possibility for TDP-43 (Afroz et al., 2017; Qin et al., 2014; Wang et al., 2018), the current study provides the first direct evidence that the NTD engages RNA, indicating that DIX domains may harbor complexities beyond mere protein-protein interactions. Notably, the interaction of NTD with RNA appears to preferentially favor GU-rich sequences, thereby sharing the specificity of the RRMs of TDP-43 (Lukavsky et al., 2013; Polymenidou et al., 2011; Tollervey et al., 2011). It is tempting to speculate that this binding may be cooperative with the canonical RRM-mediated binding and may potentially support monomeric TDP-43 interactions with large stretches of GU-rich RNA.

We showed that NTD-mediated RNA binding occurs only in the monomeric state. Structural mapping further indicated that RNA binding and oligomerization are mutually exclusive processes. This is supported by the observation that more than half of the residues involved in RNA binding are also essential for oligomerization, that both processes occupy the same structural interface, and that they rely on similar electrostatic interactions. A single phosphomimetic substitution at an RNA-interacting residue (S48E), but not a charge-neutral mutation (S48A), was previously shown to be sufficient for disrupting oligomerization *in vitro* (Wang et al., 2018), indicating that even a single negative charge at this interface can prevent NTD–NTD interactions. Therefore, we hypothesize that RNA binding at the same region would exert an even stronger effect, potentially driving monomerization of TDP-43. Further studies will decipher the conditions that favor each state, and the cellular factors that potentially regulate the balance between RNA binding and oligomerization.

We and others have extensively investigated the role of TDP-43 oligomerization in its function and pathobiology. Oligomerization is critical for incorporation of TDP-43 into biomolecular condensates (Hallegger et al., 2021; Koehler et al., 2022; Pérez-Berlanga et al., 2023; Wang et al., 2018). Disruption of TDP-43 oligomerization induces splicing dysregulation including within its own transcript, leading to disruption of the autoregulatory feedback loop that normally maintains the steady levels of the protein (Afroz et al., 2017; Jiang et al., 2017; Koehler et al., 2022; Pérez-Berlanga et al., 2023; Wang et al., 2018; Yang et al., 2025; Zhang et al., 2013). Oligomerization is also required for efficient nuclear retention and was shown to counteract pathological aggregation (Afroz et al., 2017; Duan et al., 2021; Koehler et al., 2022; Pérez-Berlanga et al., 2023), potentially by spatially separating the LCDs (Afroz et al., 2017). Consistent with this, impaired dimerization/multimerization and an abnormal increase in monomeric species has been observed in ALS postmortem tissue (Oiwa et al., 2023).

While all this extensive work has established the importance of TDP-43 oligomerization, it is perhaps surprising that a potential contribution of NTD-mediated RNA binding in the functionality of TDP-43 was not previously recognized. The reason for this oversight might be that the two modalities, NTD-NTD oligomerization and NTD-RNA binding are intertwined, as we showed in the current study. As a result, most approaches used to disrupt oligomerization, including the point mutations 6M, 5M, and 4M (Afroz et al., 2017; Oiwa et al., 2023; Pérez-Berlanga et al., 2023) and the complete NTD deletion (French et al., 2019; Grese et al., 2021; Oiwa et al., 2023; Zhang et al., 2013) simultaneously eliminate both oligomerization and NTD-mediated RNA binding. Similarly, other variants such as the phosphomimetic S48E, C39S/C50S, and G53D (Jiang et al., 2017; Wang et al., 2018; Yu et al., 2021), which are located within the NTD head region are likely to impair RNA binding, with substitutions to negatively charged residues (E, D) potentially exacerbating this effect through electrostatic repulsion.

So what is the functional role of this RNA-binding via the NTD of TDP-43? While our data demonstrate that NTD-mediated RNA binding is not required for splicing regulation or cryptic exon repression, we have not investigated other known functions of TDP-43. For example, recent studies have reported a key role of TDP-43 in alternative polyadenylation (Arnold et al, 2025; Bryce-Smith et al, 2025; Zeng et al, 2025), a function that is critical for neuronal health. Since some of these sites are characterized by long GU-rich stretches, we cannot exclude the possibility that NTD-RNA interactions support the polyadenylation site selection by TDP-43. This might be particularly relevant for cell types with intrinsically high levels of monomeric TDP-43 in the nucleus, which might need cooperation between NTD-mediated and canonical RRM-mediated binding to stabilize interactions with RNA. Intriguingly, while in most cells, TDP-43 predominantly exists in oligomeric assemblies in the nucleus (Afroz et al., 2017; Oiwa et al., 2023; Pérez-Berlanga et al., 2023; Yu et al., 2021), human motor neurons were shown to have intrinsically higher levels of monomeric TDP-43 in the nucleus compared with non-neuronal cells, both in culture and in human tissue (Huang et al., 2024). In combination with our current findings, this observation raises the possibility that NTD-mediated RNA binding may be particularly important in this cell type. Our observation that NTD-mediated RNA binding can partially support RNA-binding of full length TDP-43, even when the canonical RRM-dependent binding is impaired, is in line with this hypothesis.

Another attractive hypothesis is that the NTD-mediated RNA binding may be particularly relevant for the cytoplasmic functions of TDP-43, which remain incompletely understood. Indeed, while we and others have reported the presence of dimeric TDP-43 species in the cytoplasm (Afroz et al., 2017; Oiwa et al., 2023; Pérez-Berlanga et al., 2023), the main TDP-43 pool is likely predominantly monomeric in this compartment. TDP-43 was shown to regulate multiple aspects of post-transcriptional gene expression in the cytoplasm, including mRNA stability, transport, and translation (Birsa et al., 2020). In particular, it binds to 3′ untranslated regions (3′-UTRs) of target mRNAs and modulates their stability, either positively (e.g., *NFL*, *ADD2*) or negatively (e.g., *VEGFA*, *GRN*, *NPTX2*), thereby influencing protein expression levels (Birsa et al., 2020; Colombrita et al., 2012; Costessi et al., 2014; Hruska-Plochan et al., 2024; McDonald et al., 2011; Sidibé et al., 2021; Zhang et al., 2013). In addition, TDP-43 can directly regulate translation by binding to 3′- or 5′-UTRs, enhancing translation of specific targets such as *CAMTA1*, *MIG12*, mitochondrial transcripts (*ND3*, *ND6*), and *DENND4A* (Majumder et al., 2016; Neelagandan et al., 2019; Wang et al., 2016). Notably, several of these targets have been linked to neurodegeneration, suggesting that dysregulation of these processes may contribute to disease (Birsa et al., 2020; Hruska-Plochan et al., 2024; Neelagandan et al., 2019; Wang et al., 2016; Zhang et al., 2013). Given the critical role of the NTD and oligomerization for the phase separation of TDP-43 (Pérez-Berlanga et al., 2023; Wang et al., 2018), it is tempting to speculate that this novel NTD-RNA interaction may facilitate the formation of biomolecular condensates, which were shown to be necessary for splicing regulation (Hallegger et al., 2021; Koehler et al., 2022; Pérez-Berlanga et al., 2023). This same role may enable the incorporation of TDP-43 into phase separated compartments either in the nucleus under conditions of reduced RNA binding via the RRMs, or in the cytoplasm, where the availability of RNA targets is limited. Identifying the specific targets and (sub)cellular environments where this non-canonical binding is functional will be essential for understanding the broader regulatory roles of TDP-43 in health and disease.

While the identification of this novel RNA-binding mode is intriguing, the functional role of the monomeric NTD-RNA-bound state, and how it interplays with canonical RRM-mediated RNA binding, remains to be elucidated. Yet, this work reinforces the view that TDP-43 exists in multiple distinct conformational states, each associated with specific and perhaps unique functional entities. A deeper understanding of how these states are regulated and transition between physiological and pathological conditions will be essential for elucidating TDP-43-mediated disease mechanisms.

## Material and Methods

### Cloning

All TDP-43 constructs contained human TDP-43 sequences. Full-length TDP-43 pJ4M plasmid was acquired at addgene (Plasmid number #104480). All constructs containing individual domains were cloned into the pJ4M through deletion cloning using the Nebuilder HIFI RNA assembly cloning kit (New England Biolabs). The previously reported pLVX-EF-1α-TDP-43-HA-3′UTR construct was used as the backbone for generation of a lentiviral GFP– TDP-43 expression vector (Hruska-Plochan et al., 2024). An N-terminal eGFP tag was inserted upstream of TDP-43 using the NEBuilder HiFi DNA Assembly Cloning Kit (NEB). The two coding sequences were connected by a 42 bp linker sequence (5′-TCCGGAGGGGATCGGTGGAGCTCCACCGGTGGCTCTTCTGGC-3′).

The 6M, Tm, and Hm variants were subsequently generated in the pLVX-GFP-TDP-43 construct using the same cloning strategy. Mutant TDP-43 sequences were derived from previously described plasmids (Pérez-Berlanga et al., 2023) or generated by site-directed mutagenesis PCR using the high-fidelity Phusion DNA Polymerase (New England Biolabs) and subsequent Dpn1 (New England Biolabs) digestion before transformation. Primers and annealing temperatures used for cloning are found in Supplementary Table S1.

For CONA assays, a GFP-tagged TDP-43 expression plasmid previously described was used (Huang et al., 2024). Mutant constructs carrying the 3M (E14A/E17A/E21A), RRM1m (F147A/F149A), and combined 3M/RRM1m mutations were custom synthesized by VectorBuilder.

### Protein production, purification, and labeling

Recombinant protein production was done in One Shot™ BL21 Star™ (DE3) Chemically Competent *E. coli* (Thermo Fisher Scientific). For non-labeled protein expression bacteria were grown in Luria-Bertani (LB) broth base (Invitrogen) and for bacterial growth used for NMR experiments media was substituted with M9 minimal media supplemented with ^15^NH_4_Cl (Sigma-Aldrich) and/or D-Glucose-^13^C_6_ (Sigma-Aldrich). Protein concentrations after purification were determined with the respective extinction coefficients and molecular weights of the constructs (determined by ExPASy ProtParam, https://doi.org/10.1385/1592598900) and absorbance at 280 nm (NanoDrop, Thermo Scientific). The purity of each construct was assessed with silver or Coomassie staining.

NTD-His_6_ constructs were induced at an OD_600_ = 1.0 with 1 mM isopropyl β-D-thiogalactoside (IPTG, Biosolve) and incubated for 30 h at 25 °C. Harvested cells were resuspended in 20 ml lysis buffer (50 mM sodium phosphate (Sigma) buffer pH 8.0, 300 mM sodium chloride (NaCl, Sigma-Aldrich), 10 mM imidazole (Sigma-Aldrich), 1 mM 1,4-dithiothreitol (DTT, Thermo Fisher Scientific) and 2 protease inhibitor tablets (Roche). To lyse the cells we used a rod sonicator (W-385 LSL Secfroid sonicator, heat system ultrasonic) with 11 cycles (10 s on and 20 s off). The cell lysate was centrifuged (15,000 g, 4 °C, 20 min) and the supernatant was applied to Ni Sepharose Excel material (Sigma) in a gravity column (Econo glass column, Biorad). The column was washed once with wash buffer 1 (50 mM sodium phosphate buffer pH 8.0, 1 M NaCl, 10 mM imidazole, 1 mM DTT) and a second time with wash buffer 2 (50 mM sodium phosphate buffer pH 8.0, 1 M NaCl, 30 mM imidazole, 1 mM DTT). The protein was eluted in 30 ml elution buffer (50 mM sodium phosphate buffer pH 8.0, 1 M NaCl, 500 mM imidazole, 1 mM DTT) and dialyzed overnight (ON) in dialysis buffer (5 mM sodium phosphate buffer pH 7.4, 1 mM NaCl, 1 mM DTT). Pure protein was concentrated using Vivaspin 5,000 MWCO concentrators (Sartorius Stedium Biotech), aliquoted, flash frozen, and stored at - 20 °C.

The RRMs-His_6_ constructs were induced with 1 mM IPTG upon an OD_600_ of 0.8. The protein was expressed for 14 h at 16 °C. Harvested cells were resuspended in 30 ml/L lysis buffer (20 mM HEPES (Sigma), pH 7.5, 1 M NaCl, 10 % glycerol (Sigma-Aldrich), 30 mM imidazole, 5 mM tris(2-carboxyethyl)phosphine (TCEP, Sigma-Aldrich). The bacterial cells were lysed using a rod sonicator (W-385 LSL Secfroid sonicator, heat system ultrasonic) with 11 cycles (10 s on and 20 s off). The protein was loaded on Ni Sepharose Excel material in a gravity column, which was pre-equilibrated in equilibration buffer (20 mM HEPES, pH 7.5, 150 mM NaCl, 10 % glycerol, 5 mM TCEP). The column was subsequently washed with lysis buffer and eluted in 30 ml elution buffer (20 mM HEPES, pH 7.5, 150 mM NaCl, 10 % glycerol, 500 mM imidazole, 5 mM TCEP). The protein was dialyzed ON in dialysis buffer (20 mM HEPES, pH 7.5, 150 mM NaCl, 5 mM TCEP) concentrated using pre-equilibrated 5,000 MWCO Vivaspin 20-mL concentrators (Sartorius). The protein was subsequently aliquoted, flash-frozen, and kept at -20 °C.

NTD-RRMs-His_6_ constructs expression was done after reaching an OD_600_ of 0.7 and induced with 1 mM IPTG and subsequently incubated for 24 h at 21 °C. Cell pellets were resuspended 100 ml/L lysis buffer (50 mM HEPES, pH 7.5, 100 mM potassium chloride (KCl, Sigma-Aldrich), 10 mM MgCl_2_, 10 mM CaCl_2_, 10 mM imidazole, 10 % (v/v) glycerol, 0.5 % (v/v) Triton X-100 (Sigma-Aldrich)). Lysis was achieved using chemical lysis with 20 μg/ml DNAse, 20 μg/ml RNAse A, 1 mg/ml lysozyme, 5 protease inhibitor tablets, 5 mM TCEP) by incubating the resuspension at room temperature (RT) for 1.5 h. Lysate was cleared and the supernatant was loaded onto Ni Sepharose Excel material self-packed gravity column pre-equilibrated in column equilibration buffer (30 mM Tris, pH 8.0, 150 mM NaCl, 10 % (v/v) glycerol and 1 mM DTT). The column was subsequently washed with a wash buffer (30 mM Tris pH 8.0, 1 M NaCl, 10 % (v/v) glycerol, 1 mM DTT). Elution was achieved with 30 ml of elution buffer (30 mM Tris pH 8.0, 150 mM NaCl, 10 % (v/v) glycerol, 300 mM imidazole, 1 mM DTT) and subsequently dialyzed ON at 4 °C in dialysis buffer (20 mM potassium phosphate pH 8.0, 30 mM KCl, 10 % (v/v) glycerol, 1 mM DTT). The protein was subsequently aliquoted, flash-frozen, and kept at -20 °C.

### Sodium dodecyl-sulfate polyacrylamide gel electrophoresis (SDS-PAGE)

The same amounts of recombinant protein were mixed with LDS loading buffer (Invitrogen) and reducing agent (Life Technologies) and incubated for 10 min at 70 °C. Samples were loaded onto 15- or 17-well BOLT 4-12 % Bis-Tris precast gels (Invitrogen). Gels were run in MES buffer (Invitrogen) at 80-150 V until the lowest marker band (Thermo Scientific) reached the bottom of the gel. For purity analysis gels were incubated in Coomassie staining (Simple blue safe stain, Thermo Fisher Scientific) for 1 h at RT and subsequently washed in water for 1 h and then ON.

### CLIR-MS experiments

Briefly, a recombinant protein is incubated with an RNA, and both are covalently linked by UV radiation. The cross-linked complexes are subjected to enzymatic digestion with both RNases and proteases, and peptide-RNA conjugates are enriched using titanium oxide affinity chromatography. Enriched RNA-peptide conjugates are labeled with an equimolar mixture of ATP containing the natural oxygen isotope (^16^O-ATP) or heavy isotope-labeled ATP (^18^O-ATP), resulting in conjugates with a distinct mass difference (Δm) of 6 Da due to the heavy and light labeling. Peptides cross-linked to labeled RNA show a specific isotopic pattern with the given Δm in the precursor mass scans (MS1). Products are then fragmented, and tandem mass spectra (MS2) are used to localize the nucleic acid cross-link position in the peptide backbone by using the isotopic shift when comparing the light and heavy MS2 spectra, thus allowing for precise mapping of RNA interaction sites.

Recombinant protein-RNA cross-linking was performed in 5 mM sodium phosphate, pH 7.4, 1 mM NaCl, 1 mM DTT. NTD-RRMs-His_6_ or RRMs-His_6_ with (GU)_6_ or (AC)_6_ RNA (Microsynth AG, Switzerland) were irradiated at 50 µM final concentration and a total of 200 µg in the presence of 50 µM RNA after being incubated on ice for at least 10 minutes. The sample was split into 20 µL aliquots in a 20-well Terasaki plate (Sarstedt) and irradiated with 0.4 J/cm^2^. Mutant RRMs-His_6_ (RRM^mut^) was prepared identically with the sole difference of an irradiation strength of 3.2 J/cm^2^. Cross-linking of NTD-His_6_ for discovery purposes with (GU)_6_ and (AC)_6_ was done at 50 µM final concentration and a total protein amount of 75 µg in the presence of 50 µM RNA. For quantitative measurements, NTD^WT^ and NTD^Tm^ were cross-linked at a concentration of 50 µM final concentration and a total of 100 µg in the presence of 50 µM (GU)_6_ RNA. Samples were incubated on ice for 30 min and subsequently split into 20 µL aliquots in a 20-well Terasaki plate and irradiated with 3.2 J/cm^2^. The experiment was performed in quadruplicates for both protein variants. For irradiation with 254 nm UV light, samples were placed on ice at a distance of 3 cm from the lamps of a Spectralinker XL-1500 device (Spectronics Corp.).

Sample processing followed previously established procedures(Dorn et al., 2017; Götze et al., 2021; Sarnowski et al., 2023). In brief, after cross-linking, samples were precipitated, then digested with a mixture of RNases A and T1 (at 52 °C for 2 h), followed by protease digestion using trypsin (at 37 °C ON) and a second round of RNase A/T1 treatment. Digested samples were purified by solid-phase extraction (SPE) and labeled with a 1:1 mixture of ^16^O-ATP and ^18^O-ATP, followed by a second SPE step. Peptide-RNA conjugates were enriched by titanium dioxide affinity chromatography and purified by miniaturized SPE in StageTips. Samples were analyzed on an Easy nLC-1200 HPLC system coupled to an Orbitrap Fusion Lumos mass spectrometer equipped with a Nanospray Flex ion source (all ThermoFisher Scientific). Analytes were separated on an Acclaim PepMap RSLC C18 column (250 mm × 75 µm i.d., 2 µm particle size, ThermoFisher Scientific) at a flow rate of 300 nL/min and with mobile phases A = water/acetonitrile/formic acid (98:2:0.15, v/v/v) and B = acetonitrile/water/formic acid (80:20:0.15, v/v/v). A linear gradient from 6 to 32 %B in 60 min was used for the separation. For identification of cross-linking sites by data-dependent acquisition (DDA), samples were dissolved in 15 µL of water/acetonitrile/formic acid (95:5:0.1, v/v/v) and duplicate injections of 5 µL were performed. DDA was performed in top-speed mode with a cycle time of 3 s. Precursors were detected in an orbitrap full scan at a resolution of 120,000 and fragment ions with a charge state between 2+ and 7+ were selected for HCD fragmentation using a stepped normalized collision energy of 23 ± 5 %. Fragment ions were detected in the linear ion trap (rapid scan mode). Dynamic exclusion was activated for 30 s after one sequencing event.

For quantitative parallel reaction monitoring (PRM) all samples were dissolved in 15 µL of water/acetonitrile/formic acid (95:5:0.1, v/v/v) and single injections of 3 µL were performed. The PRM method was set up with a precursor full scan in the orbitrap analyzer at a resolution of 60,000 and a series of targeted MS/MS events using HCD fragmentation with a stepped normalized collision energy of 23 ± 5 %, as above. Fragment ions were detected in the orbitrap at a resolution of 60,000 with an AGC target of 4.0e5 and a maximum injection time of 200 ms. Two PRM target lists were used, with each list containing 10 m/z ratios (five different peptide-RNA conjugates in two charge states, 2+ and 3+) and with targets resulting from ^16^O-labeling monitored in one run and targets resulting from ^18^O-labeling in a second run. No retention time scheduling was used because of the broad elution range of the targets.

For data analysis of the identification of cross-linking sites by data-dependent acquisition (DDA), cross-linking sites were identified using RNxQuest, version 1.0 (Sarnowski et al., 2023), based on xQuest, version 2.1.5 (Rinner et al., 2008; Walzthoeni et al., 2012). Mass spectrometry data in the Thermo .raw format were converted into the .mzXML format using msconvert/ProteoWizard, version 3.0.9393 (Chambers et al., 2012), and searched against the sequence of the target construct and its reversed sequence, and the corresponding RNA sequences. The maximum length for an RNA adduct was set to four nucleotides for all experiments. Oxidation of methionine was set as a variable protein modification, for RNA adducts the following product types were considered for all experiments: unmodified, -H_2_ (2.01565 Da), -H_2_O (18.010565 Da), -HPO_3_(79.966331 Da), - HPO_3_+H_2_O (61.955766 Da), - H_3_PO_3_ (81.981981 Da), MS/MS spectra were paired according to the distinct mass shift of 6.012735 Da resulting from ATP labeling. Additional search settings were: enzyme = trypsin, the maximum number of missed cleavages = 2, precursor ion tolerance = 10 ppm, fragment ion tolerance = 0.2 Da for “common” fragment ions (not containing the RNA modification), and 0.3 for “xlink” ions (containing the RNA modification). The false discovery rate was set to 1 % at the peptide-RNA conjugate level.

For the analysis of changes in RNA binding propensity by the NTD^WT^ and NTD^Tm^ PRM transition lists containing all possible “common” and “xlink” fragment ion masses were imported into Skyline software (64-bit, 22.2.0.351) (MacLean et al., 2010). Precursor ions were manually identified based on the coelution of light and heavy fragment ions, matching the intact peak shape. Peak boundaries were adjusted to exclude interfering and/or closely eluting peaks. The abundance of targeted RNA adducts was determined by summing up the integrated peak areas of the six most intense fragment ions per precursor, in both light and heavy channels. Peptide intensity values in different samples were normalized by the intensity of two non-crosslinked TDP-43 NTD peptides that co-purified. Identified outliers were excluded based on normality (Shapiro-Wilk test) and deviating standard deviations (Brown-Forsythe test).

### Molecular dynamics (MD) simulations

Three molecular systems were investigated by molecular dynamics: a TDP-43 NTD monomer system with the oligoribonucleotide GUGU, a monomer system with ACAC, and a dimer system with GUGU. The set-up was primarily based on the structure 5mdi (T. Afroz et al., 2017). For the two monomer systems, chain A of 5mdi was used, which had a protein sequence ranging from 1 to 79. For the dimer system, two chains of 5mdi were used, and the two NTDs had the same protein sequence (1-79). 3D coordinates of GUGU were extracted from structure 4BS2 (Lukavsky et al., 2013) and ACAC was built by mutating the corresponding side chains of GUGU. To ensure sufficient separation between NTD and oligoribonucleotides, their centers of mass were kept at 25 Å. Crystal water molecules were retained in the built systems, and hydrogen atoms were added using the CHARMM program (version 45b2) (Brooks et al., 2009). We determined protonation states according to hydrogen-bond networks at neutral pH conditions by Maestro (12.8, Schrödinger, LLC: New York, NY, 2021). The complex system was solved in a rhombic dodecahedron (RHDO) TIP3P (Jorgensen et al., 1983) water box (lattice length: 91 Å for the two monomer systems and 95 Å for the dimer system), ensuring at least a 12 Å buffer distance between the system atoms and the boundary of the water box. Finally, Na+ and Cl– ions were added to the solvated systems to neutralize them and mimic physiological conditions, at a concentration of 0.15 M. To release poor contacts and geometries, each simulation system was initially minimized for 10,000 steps under a series of restraints and constraints on the solute atoms. The minimized structure was subsequently heated to 300 K and equilibrated in an NVT condition (constant volume and temperature) and in a followed NPT condition (constant pressure and temperature). All equilibration phases lasted 10 ns and were conducted using the CHARMM program (version 45b2) (Brooks et al., 2009).

A production of MD simulation was conducted for 200 ns in the NPT condition for each trajectory using the NAMD program (version 2.13) (Phillips et al., 2005). Each system was replicated 10 times. The pressure was controlled using the Nosé–Hoover Langevin piston method with a 200 ps piston period and 100 ps piston decay time (Feller et al., 1995; Martyna et al., 1994), and the temperature was maintained at 300 K using the Langevin thermostat with a 5 ps friction coefficient. The integration time step were kept to 2 fs by constraining all bonds involving hydrogen atoms with the SHAKE algorithm. van der Waals (vdW) energies were calculated using a switching function with a switching distance of 10-12 Å (Steinbach and Brooks, 1994), and electrostatic interactions were evaluated using the particle mesh Ewald summation (PME) method (Essmann et al., 1995). Additionally Lennard-Jones long-range correction was enabled (Shirts et al., 2007).

To describe the protein and oligoribonucleotide molecules, the CHARMM36m force field was used (Huang et al., 2013). 10 independent runs were performed for each system with random initial velocities, totaling in 2 μs of simulation time. Snapshots of the MD trajectories were saved every 20 ps for further analysis. CHARMM routines were used, specifically COOR and DMAT, to analyze the contact map by counting the interaction frequency between NTD and oligoribonucleotides.

### Nuclear Magnetic Resonance (NMR) spectroscopy

To acquire NMR spectra of the NTD buffer was exchanged to NMR buffer (5 mM sodium phosphate buffer pH 7.4, 1 mM NaCl, 1 mM 2-mercaptoethanol). The Bruker AVIII-700 MHz was utilized for all spectral acquisition. To process the data the TopSpin4.0.7 (Bruker) software was used. For analysis and peak assignment, Sparky 3.133 (http://www.cgl.ucsf.edu/home/sparky) was used. For the backbone assignment of the TDP-43 NTD^Tm^ construct the following NMR experiments were measured (in H_2_O): 2D ^15^N–^1^H HSQC, 3D HNCA, 3D CBCACONH, and 3D HNCACB. All spectra were recorded at 313 K. Chemical shift perturbations were performed at 100 µM protein concentration and with increasing RNA concentrations. The chemical shift perturbations were calculated at saturation, at a protein:RNA ratio of 1:8 with the formula (Δδ = ((δHN)^2^ + (δN/6.51)^2^)^1/2^). RNA used in these experiments were ordered from Microsynth AG, Switzerland.

### Isothermal titration calorimetry (ITC)

ITC experiments were performed on a PEAQ-ITC instrument (Microcal), calibrated according to the manufacturer’s instructions. Protein and RNA samples were dialyzed against the NMR buffer. Concentrations of proteins and RNAs were determined using optical density absorbance at 280 nm and 260 nm, respectively. 20 μM of protein was titrated with 2.25 mM of RNA using 19 injections of 2 μl every 2.5 min at 30 °C. Raw data were integrated, normalized for the molar concentration, and analyzed using the MicroCAL PEAQ-ITC software of Malvern. All measurements were performed in triplicates. RNA used in these experiments were ordered from Microsynth AG, Switzerland.

### Structural image representation and movie

We generated structural figures and movies using PyMOL graphic software (Version 2.3).

### CONA assay

Experiments were done as previously described (Huang et al., 2024). In brief: Transfected cells grown in 35-mm dishes were lysed in 250 μL PBS containing 1% Triton X-100 and 40 U/mL RiboLock for 15 min on ice with intermittent vortexing. Lysates were clarified by centrifugation at 13,000 rpm for 15 min at 4°C and stored at −80°C until use.

HPLC-grade (GU)₁₅ RNA oligonucleotides labelled with Cy5 at the 5′ end and biotin-TEG at the 3′ end were custom synthesized by Eurofins. Ni-NTA Superflow beads (Qiagen, Cat. No. 30410) were washed with binding buffer (0.3 M NaCl, 20 mM HEPES pH 7.5, 0.01% Triton X-100, 5 mM MgCl₂) and resuspended as a 50% slurry. Twenty-five microlitres of bead slurry were incubated with 40 pmol His-tagged streptavidin (NKMAX, STV0801) for 30 min at room temperature with shaking. Beads were washed three times with binding buffer, resuspended, and incubated with 50 pmol RNA oligonucleotide for 30 min at room temperature with shaking. Following three additional washes with binding buffer, 5 μL of resuspended beads were added to thawed cell lysates and incubated for 2 h at room temperature with rotation. Lysate input was normalized to ensure equal amounts of GFP-tagged protein, as determined by western blot analysis.

Following incubation, beads were washed three times with binding buffer and dispensed into µCLEAR 384-well plates (Greiner, 781092) at 30 μL per well. Imaging was performed using an Opera Phenix high-content screening system (PerkinElmer). Acquisition settings were as follows: brightfield, 20 ms exposure, 50% laser power, 2.0 μm focus offset; Cy5, 100 ms exposure, 50% laser power, 2.0 μm focus offset; GFP, 100 ms exposure, 20% laser power, 2.0 μm focus offset. Z-stacks were acquired from 16.0 to 17.0 μm using three planes spaced 0.5 μm apart. Beads located both at the centre and periphery of each well were included in the analysis.

Mean ring fluorescence intensities in the Cy5 and GFP channels were quantified using a custom ring-segmentation pipeline implemented in Harmony v4.9 software (details available upon request).

Plasmid 3M (#237544) and RRM1’ (#237545) were deposited on addgene. Sequences of additional variants are to be found in Supplementary table 2.

### Lentiviral vector production

The lentiviral transfer vector was generated, harvested, and concentrated as previously described (Hruska-Plochan et al., 2024). Lentiviral pellets were resuspended in PBS to generate 10× concentrated viral preparations. To determine the appropriate transduction dose, viral titration experiments were performed in HeLa TDP-43 KO cells using increasing volumes of concentrated viral suspension.

### GFP-TDP-43 HeLa KO reporter line generation

HeLa TDP-43 knockout (KO) cells, kindly provided by Shawn M. Ferguson (Roczniak-Ferguson and Ferguson, 2019), were maintained at 37°C in a humidified incubator with 5% CO₂. Cells were cultured in DMEM/F-12 (Sigma-Aldrich, D5671) supplemented with 1× GlutaMAX (Gibco, 35050-061), 10% fetal bovine serum (FBS; Gibco, A5256701), and 1% penicillin–streptomycin (Gibco, 15140122).

A polyclonal HeLa cell line was generated by co-transfecting the piggyBac TDP-REG construct together with a plasmid encoding a hyperactive piggyBac transposase at a 1:1 ratio. Both the TDP-REG expression vector (Wilkins et al., 2024) and the hyperactive transposase construct(Yusa et al., 2011) have been described previously. Transfections were performed in 6-well plates (TPP) using HeLa TDP-43 KO cells and their parental (wild-type) counterpart at approximately 90% confluency. Lipofectamine 3000 (Thermo Fisher Scientific) was used for transfection, with 5 µL Lipofectamine and 5 µL P3000 reagent per well diluted in Opti-MEM (Gibco), following the manufacturer’s instructions.

Stable integrants were selected by culturing cells in medium containing 10 µg/mL blasticidin (InvivoGen, ant-bl-05) for two weeks. A parallel control transfection lacking the transposase plasmid was included to ensure complete elimination of transiently transfected cells during selection.

For generation of a monoclonal line, cells were sorted as single cells into individual wells of µ-Plate 96-well square ibiTreat plates (IBIDI) pre-filled with growth medium, using a BD FACSymphony S6 5L flow cytometer (BD Biosciences). Cell preparation followed the same procedure as in other FACS experiments. After approximately two weeks, a single clone was selected and subsequently maintained as a stable monoclonal line under continuous selection with 10 µg/mL blasticidin.

### GFP-TDP-43 transduction and FACS sorting

TDP-REG HeLa cells (TDP-43 knockout and wild type) were seeded at a density of 75,000 cells per well in 6-well plates. The following day, cells were transduced with lentiviral constructs encoding different GFP–TDP-43 variants. After 24 hours, the medium was replaced to remove remaining viral particles. Cells were then maintained for six days post-transduction, including a single passage into 10 cm dishes during this period.

For flow cytometry and fluorescence-activated cell sorting (FACS), cells were harvested by trypsinization and resuspended in FACS buffer composed of 1% FBS (Gibco), 25 mM HEPES (Sigma-Aldrich), and 5 mM EDTA (Sigma-Aldrich) in 1× PBS (Gibco). The suspension was passed through a 35 µm cell strainer-capped Falcon tube (Corning) to ensure a single-cell preparation prior to analysis on either a BD FACSymphony or BD FACSymphony S6 5L flow cytometer (BD Biosciences).

Cells were sorted based on two defined GFP intensity ranges. To maintain consistency of gating across experiments, GFP calibration beads (Thermo Fisher Scientific) were used as a reference. Sorted populations were collected into 1.5 mL tubes containing HeLa growth medium and kept on ice.

Flow cytometry data were analyzed using FlowJo (v10.10.0). Debris was removed by gating on forward scatter area (FSC-A) versus side scatter area (SSC-A). Doublets were subsequently excluded based on FSC-A versus FSC-H (height). The remaining singlet population was used for downstream analysis.

### Fluorescence microscopy and analysis of HeLa TDP-43 KO reporter lines

Sorted cells were seeded into µ-Plate 96-well square ibiTreat plates (IBIDI) at a density of 15,000 cells per well. The following day, cells were fixed with formaldehyde (Thermo Fisher Scientific) at a final concentration of 4% for 20 minutes at room temperature. After three washes with PBS (Gibco), samples were stored at 4°C until further use. Cells were stained with 1:1000 DAPI for 20 min at RT, washed three times in PBS. Confocal imaging was performed using a Leica SP8 inverse Falcon microscope as well as an MD ImageXpress Confocal HT.ai high-content imaging system (Molecular Devices), both operated at the Center for Microscopy and Image Analysis (ZMB) at the University of Zurich.

Nuclei were segmented using the StarDist 2D plugin in ImageJ. Fluorescence intensities for mScarlet and GFP were quantified within the segmented regions, and all subsequent analyses were performed in RStudio. To remove debris, objects with DAPI intensities exceeding 1’500 or areas smaller than 2’500 pixels were excluded.

To ensure comparable GFP signal levels across conditions, cells with GFP intensities above 700 were discarded, based on the distribution observed in the 6M condition, which exhibited the lowest overall GFP intensity. Finally, datasets were equalized by random downsampling to 44–54 cells per condition prior to downstream analysis.

### RNA sequencing and analysis

For RNA sequencing, cells were collected by sorting into 1.5 mL tubes containing culture medium, followed by centrifugation at 300 × g for 5 minutes to pellet the cells. Pellets were rapidly frozen in liquid nitrogen and stored at −80°C until further processing.

RNA was isolated from four biological replicates per condition using the RNeasy Plus Mini Kit (Qiagen, Cat. No. 74134). Lysates were homogenized with QIAshredder columns (Qiagen, Cat. No. 79654) according to the manufacturer’s protocol. Sequencing libraries were generated using the Tecan Universal Plus mRNA-Seq kit. Samples were then sequenced on an Illumina NovaSeq X Plus system using a paired-end 150 bp setup, aiming for a total yield of approximately 500 million reads.

Fastq files obtained from the sequencing run were processed through the nf-core/rnaseq v3.25.0 pipeline via NextFlow using default settings (Ewels et al., 2020; Patel et al., 2026). Briefly, read quality was assesed using FastQC, reads were then trimmed for adapter removal using TrimGalore and remaining ribosomal RNA was removed by SortMeRNA. Reads were then aligned to the GRCh38 reference human genome using STAR and then gene level quantifications were performed with Salmon using GENCODE v42 annotation. All tool versions are reported at github (Patel et al., 2026). Differential expression analysis was performed using DESeq2 (v.1.48.2) considering as expressed only the genes with at least 1 average FPKM in one of the considered conditions. Transcripts were classified as differentially expressed when displaying an adjusted P value below 0.05 (Love et al., 2014).

For correlation analysis, gene expression was quantified for each condition as the average FPKM across replicates and standardized by computing z-scores. For the gene set of interest (defined as genes differentially expressed between non-treated (NT) and WT) pairwise comparisons across all conditions were performed by calculating Pearson correlation coefficients using the stats package in R.

Hierarchical clustering of samples was performed using variance-stabilized counts obtained via the variance-stabilizing transformation implemented in DESeq2 (Love et al., 2014). Sample-to-sample distances were computed using Euclidean distance on the transformed expression matrix, and hierarchical clustering was conducted using the complete linkage method. The resulting dendrogram was visualized using the dendextend (v.1.19.1) package (Galili, 2015).

Differential splicing analysis was performed according to Brow et al., 2022, using STAR-aligned BAMs as input to MAJIQ (v2.1) and the GRCh38 reference genome. Cryptic junction annotation was obtain from previously published work (Brown et al., 2025; Vaquero-Garcia et al., 2016). All the volcano plots were made using the EnhancedVolcano (v.1.26.0) R package. Cryptic HDGFL2 junction counts were obtained from Brown et al., 2025. Sashimi plot was made using trackplot (v.0.5.3) (Zhang et al., 2023).

### Statistical analysis

All data sets were tested for their normality (Shapiro-Wilk test) and standard deviations (Brown-Forsythe test). Statistical analysis was performed as specified in the figure legends. Statistical significance was denoted as follows: ns for not significant, * for p < 0.05, ** for p < 0.01, *** for p < 0.001, and **** for p < 0.0001. Error bars in graphs depict the standard deviation (SD) of the dataset. Notably, when data were expressed as a percentage of control, analyses were conducted on the original data to maintain the integrity of variances and normal distribution (Lew, 2007).

**Table 1:**
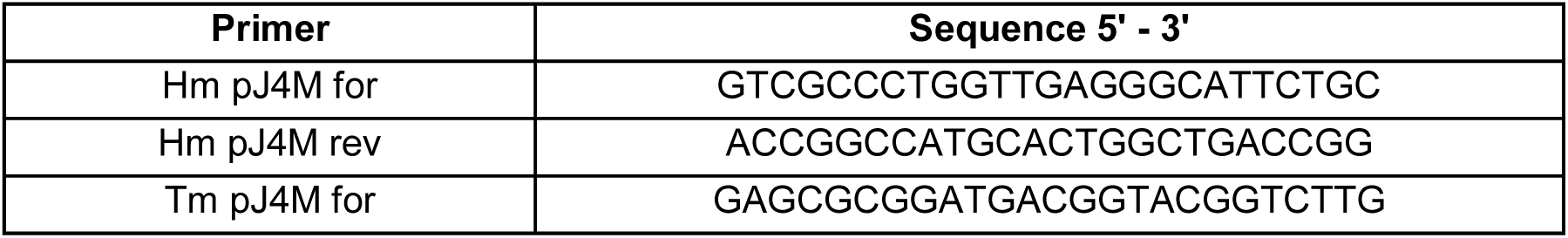
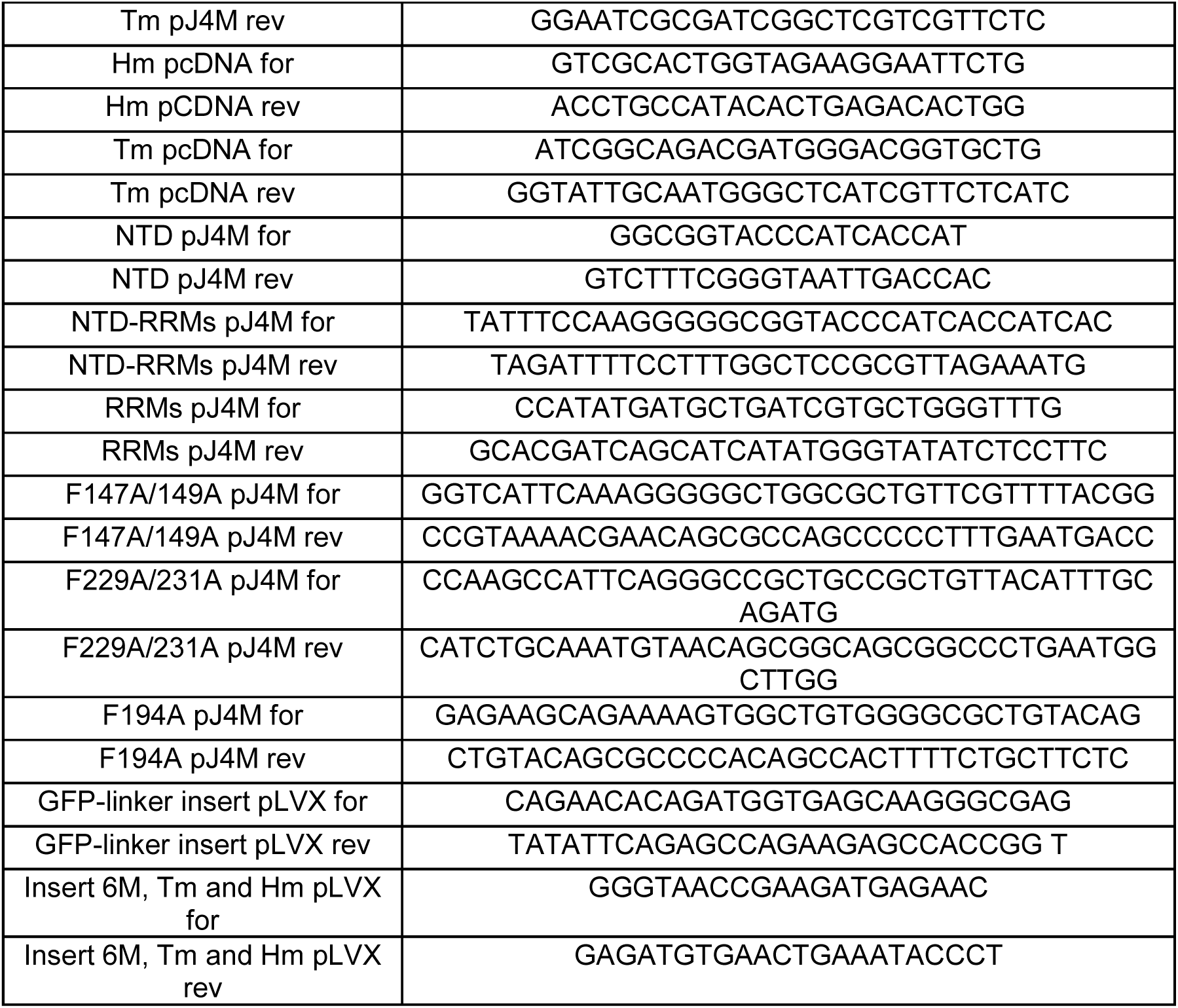
Cloning primers used in this study.

## Contributions

A.Z. led the study and conducted *in vitro* and RNA sequencing experiments, conceptualized, wrote and edited the original draft. A.O. and V.I.W. generated the reporter HeLa KO cell line and performed experiments. RNA sequencing and analysis was done by F.P. and A.Z.. I.S., M.G., N.B. and A.L. acquired and analysed the CLIR-MS data sets. Y.L. performed molecular dynamics simulations. W.H. performed and analysed the CONA assays. A.Cl. and A.Z. performed the NMR and ITC experiments. M.P., A.L., F.A., P.F.,T.S., and A.C. gave scientific advice.

## Acknowledgments/Funding

The authors would like to thank all members of the Polymenidou lab and Sinergia group for the critical feedback and discussions. A.L. would like to thank Ruedi Aebersold and Paola Picotti for access to instrumentation and laboratory infrastructure.

